# Dynamic hybrid-hydrogel lung models decouple matrix composition, stiffness, and fibroblast conditioning to define distinct drivers of fibrotic progression

**DOI:** 10.64898/2026.07.21.739843

**Authors:** Rachel Blomberg, Mikala C. Mueller, Thao Vu, Dema H. Essmaeil, David W. H. Riches, Chelsea M. Magin

## Abstract

Idiopathic pulmonary fibrosis is a devastating chronic lung disease characterized by progressive scarring of the lung, which leads to impaired gas exchange and ultimately death. While research has provided us with extensive understanding on end-stage disease, the factors that lead to forward-feedback loops of fibrotic progression are still not fully known. Cell intrinsic phenotype, pathological extracellular matrix (ECM) composition, and increased tissue stiffness are all hallmarks of advanced fibrosis, but the relative contribution of these factors to disease has been difficult to disentangle using classic *in vivo* models. In this study we created biomaterials-based 3D lung models that incorporate geometrically relevant co-culture of lung epithelial cells and fibroblasts with tunable stiffness, ECM-containing hybrid-hydrogels. Using this model system, we demonstrated that environmental stiffness has the strongest effect on overall fibroblast phenotype. RNAseq analysis revealed unique gene-level changes in both fibroblasts and epithelial cells due to both composition and stiffness, highlighting the importance of incorporating both factors into any *in vivo* disease models. Overall, these results reinforce the value of biomaterials-based models in understanding and disentangling contributors to disease pathogenesis.

## Introduction

Idiopathic pulmonary fibrosis (IPF) is the most common interstitial lung disease, characterized by progressive and persistent scarring of the lungs, resulting in impaired breathing, gas exchange, and ultimately death.^1^ Current FDA-approved treatments for IPF can only control symptoms rather than reverse the course of disease, but development of new therapeutics has been limited by the incomplete ability of current preclinical models to accurately replicate human disease.^2^ Our current understanding of the etiology of IPF is that injury to the lung epithelium initiates a regenerative response involving the differentiation of epithelial cells, but that this process can be interrupted, leaving cells in a transitional state.^3,4^ In this state, they can acquire characteristics of basal or even mesenchymal cells and trigger a pathological wound-healing response in the local environment.^3,5^ Quiescent fibroblasts are activated in response to these cues and begin to synthesize and remodel a dense, fibrillar extracellular matrix (ECM), which increases the local tissue stiffness.^6^ This stiff, pathological matrix promotes a forward-feedback loop of fibroblast activation and epithelial cell differentiation, resulting in the progressive nature of the disease.^7^

Based on these collective pathogenic features, there are several important factors to consider when modeling IPF progression *in vivo*. Cell intrinsic and extrinsic factors both clearly play a role, with coordinated changes in both ECM composition and biomechanics driving extrinsic forces. Multiple studies have assessed the phenotype of fibroblasts isolated from IPF lung and demonstrated that in traditional cell culture environments (e.g. monolayer culture on plastic), fibrotic fibroblasts retain unique phenotypes characterized by proliferation (reduced), oxidative stress response, apoptotic resistance, and expression of classic activation markers like alpha smooth muscle actin (αSMA) and collagen 1 alpha chain 1 (Col1a1).^8–10^ When fibroblasts from IPF or normal lung are cultured on collagen, these cells also demonstrated differences in proliferation mediated by integrin signaling.^11^ Critically, IPF fibroblasts cultured on plastic surfaces show reduced proliferation, but when grown on collagen, proliferation is increased, highlighting the importance of culture environment on cell behavior. Decellularized lung scaffolds, generated by treatment of lung tissue with a variety of detergents and/or enzymes, can be used to culture cells in a 3D environment that preserve the compositional, architectural, and mechanical differences present in IPF.^12^ When healthy fibroblasts are cultured on decellularized IPF scaffolds, the presence of a fibrotic matrix is sufficient to induce cellular activation.^12,13^ While these scaffolds are a powerful tool for *in vitro* studies, decellularized tissues do not allow for decoupling of the biochemical and biomechanical inputs that can drive fibrotic progression. In a far more reductionist system, fibroblasts can be cultured in 2D on polyacrylamide (PA) hydrogels coated with chosen ECM substrates. One study using this system noted that lung fibroblasts were activated in response to stiffness when cultured on fibronectin, but culture on collagen promoted activation regardless of stiffness,^14^ highlighting the potential of ECM composition and stiffness to independently impact fibrotic activation and progression. Similarly, using 2D polyacrylamide hydrogels of varying stiffness, coated with ECM derived from normal or IPF lung, Sava et. al. demonstrated that stiffness is sufficient to induce myofibroblast-like phenotypes in pericytes, regardless of ECM composition.^15^ Environmental stiffness has also been shown to overcome cell-intrinsic activation, as fibrotic fibroblasts cultured on PA hydrogels that replicate healthy lung stiffness revert to a more quiescent phenotype.^16^ Collectively, these observations highlight a critical need for engineered culture platforms that can decouple cell-intrinsic phenotypes from extracellular matrix composition and mechanics, enabling mechanistic investigation of the distinct factors that drive fibrosis.

Dimensionality is another critical consideration in designing *in vitro* culture platforms. The results demonstrated in 2D studies, as above, have not always been fully reproduced in 3D culture systems, where the ability of cells to undergo cytoskeletal remodeling for spreading or contractility can be limited by other mechanical properties of their environment.^17^ Cells in 3D may exhibit differential spreading based on the mechanical loading,^18^ stress relaxation,^19,20^ and/or degradability^21^ of their culture environment, and even where cells demonstrate an ability to spread in 3D, a commonly observed morphology involves rounded cell bodies with filamentous protrusions, rather than a fully extended cytoplasm as observed in 2D.^18,19,21^

Our lab has engineered dynamic decellularized ECM (dECM)-containing hybrid-hydrogels that enable independent control of ECM composition and matrix mechanics for mechanistic studies of fibrosis. Hybrid-hydrogels are comprised of an 8-arm poly(ethylene glycol) macromer, with the ends of each arm functionalized with a reactive alpha-methacrylate group (PEGαMA). dECM is generated by homogenizing, decellularizing, digesting, and thiolating lung tissue.^22,23^ The resultant mixture contains a spectrum of physiologically relevant proteins and glycosyaminoglycans, including structural components like fibrillar collagen and a variety of ECM-associated growth factors.^24^ Unlike conventional dECM scaffolds, which inherently couple matrix composition and mechanics, thiolation allows dECM fragments to act as multifunctional crosslinkers in a nucleophile-catalyzed thiol-ene Michael addition reaction within a synthetic PEG network. The presence of tyrosine residues on dECM fragments can subsequently be leveraged to induce on-demand dynamic stiffening through ruthenium/SPS-mediated dityrosine dimerization. These orthogonal chemical reactions allow matrix mechanics to be altered independently of dECM source, creating a unique platform that decouples biochemical and biomechanical cues while preserving physiologically relevant matrix complexity. Using these chemistries, hydrogels can be tuned to mimic the stiffness of either healthy or diseased lung, regardless of the original source of dECM. Our previous studies with hybrid-hydrogels in 2D, demonstrated that stiffness exerts a greater influence on fibroblast activation than matrix composition.^23^ Recognizing that cellular responses differ substantially between 2D and 3D environments, we designed multicellular 3D lung models composed of alveolar-like epithelial structures surrounded by fibroblast-laden hydrogels. These models revealed stiffness-mediated pro-fibrotic crosstalk between epithelial cells and fibroblasts.^5^ Here, we integrate dynamic hybrid-hydrogel engineering with multicellular 3D lung modeling, to independently manipulate fibroblast pre-conditioning, matrix composition, and tissue stiffness, enabling direct interrogation of distinct and interacting contributions to fibrotic progression.

## Methods

### Animal use and bleomycin model

All animal use was carried out in accordance with IACUC-approved protocols at the University of Colorado Anschutz (protocol #1002). To generate fibrotic murine lungs, 8–12-week-old C57BL/6 mice were administered a single dose of 3 U kg^-1^ bleomycin (MWI Veterinary Supply) via intratracheal instillation. Mice were euthanized 21 days after bleomycin administration, as previously described, at which point both inspiratory capacity and quasi-static compliance of the lungs was significantly reduced by around 40% in bleomycin-exposed animals.^23^ For all experiments, equal numbers of male and female mice were pooled to account for potential variability due to sex, since the goal of this study was not to investigate sex differences.

### PEGαMA synthesis and characterization

PEGαMA was synthesized as previously described.^22,24^ Hydroxylated PEG (0.004 mol - OH) was dissolved in anhydrous tetrahydrofuran (THF), combined with anhydrous sodium hydride (0.015 mol), and reacted at room temperature for 45 minutes. Ethyl 2-(bromomethyl)acrylate (0.026 mol) was then added dropwise to the reaction, which was allowed to proceed for 48 h, protected from light. The reaction was then neutralized with 1 N acetic acid and the product filtered through celite 545 before concentration via rotary evaporation. Product was precipitated in diethyl ether overnight at 4°C and then dialyzed in 1 kg mol^-1^ tubing for a total of three changes over three days. Product was lyophilized and functionalization confirmed via ^1^H NMR on a Bruker DPX-400 FT NMR spectrometer (300 MHz).^25^

### dECM generation and functionalization

Control or bleomycin-injured 11–15-week-old mice were euthanized by CO_2_ inhalation followed by bilateral thoracotomy. Cardiac perfusion of 20 ml phosphate-buffered saline (PBS) was performed to clear blood from the lungs, and then lungs were removed for decellularization. Lungs from 10 mice (5 male and 5 female) were pooled into 10 ml deionized (DI) water and homogenized using a GentleMACs tissue homogenizer (setting lung 2.01; repeated three times). The resulting slurry was poured into a 40 µm Uberstrainer cell strainer (Pluriselect) and washed by pipetting through 5 ml DI water three times. 5 ml of 0.2% Triton X-100 (in water) was added to the strainer, which was quickly capped and inverted for overnight incubation at 4°C. Following incubation the material was washed three times with DI water then incubated overnight at 4°C in 2% v/v sodium deoxycholate in DI water. Material was washed three times with water, incubated for 1 h at 37°C in 2M sodium chloride in DI water, washed three times with water, and then incubated 1 h at 37°C in 12 U mL^-1^ DNase. After three final water washes, decellularized material was collected into a 50 ml conical tube and lyophilized. Lyophilized material was digested with pepsin (320 U mg^-1^ in 0.1N HCL) for 36 h before being neutralized with 0.1 N NaOH and lyophilized again. Digested dECM was functionalized via thiolation by reaction with 75-molar excess Traut’s reagent for 1 h at room temperature.^23^

### Hydrogel fabrication and rheology

Hybrid-hydrogels were fabricated by reacting PEGαMA (10-11 wt%) with a crosslinker solution containing 50% dECM, 25% MMP-2 degradable peptide sequence (KCGPQGIWGQCK, GL Biochem), and 25% PEG-dithiol (average M_n_ 3,400; Sigma Aldrich) at a ratio of 0.75 thiols to -enes. dECM was first dissolved in 30 mM TCEP (pH 8.0) by 30-min incubation in a sonicating water bath. The remaining hydrogel precursors were dissolved in 2-aminomethyl-1H-imidazole dihydrochloride (10 mM, pH 8.0; Acros Organics) and then combined to form the hybrid-hydrogel precursor solution. Hydrogels were incubated at 37° for 20 minutes to ensure full polymerization via nucleophile-catalyzed Michael addition.^26^ For rheological characterization, 8-mm hydrogel discs were synthesized, swollen in PBS overnight, and analyzed on a Discovery HR2 rheometer (TA Instruments) by measuring storage modulus (G’) under a frequency oscillation with logarithmic sweep of frequencies between 1 and 100 rad s 1 at 1% strain.^22,23^ Data are presented as storage modulus of 3-4 individually synthesized batches of hydrogels, with n = 3 individual hydrogels assessed per batch.

### 3D lung model formation and culture

3D lung models were generated as previously described.^5^ Briefly, hydrogel microspheres comprised of a PEG-norbornene macromer, MMP3-degradable peptide crosslinker, and adhesion peptides CGRGDS and CGYISGR were generated by emulsion polymerization.^5^ Primary murine epithelial cells were isolated on a magnetic column by binding to anti-EpCAM magnetic beads, resulting in a population that is approximately 80% SFTPC+ alveolar epithelial type 2 cells.^5^ Epithelial cells were labeled with magnetic nanoparticles (Nanoshuttle; Greiner Bio-One) and mixed with hydrogel microspheres (∼200 µm diameter) at a ratio of 250 cells/sphere under a magnetic field and on an orbital shaker for three days.^5^ On the third day, primary murine alveolar fibroblasts were isolated using anti-PDGRFα magnetic beads and suspended in hybrid-hydrogel solution. For fibroblast imaging experiments, cells were isolated from dual reporter αSMA.RFP/Col1a1.GFP mice.^27^ Epithelial cell and microsphere aggregates in a multi-well plate were transferred to a concentrating magnet placed below the plate while media was carefully removed. Aggregates were ringed with a hydrophobic pen (ImmEdge) and fibroblast-containing hybrid-hydrogel solution was pipetted on top to encapsulate. Plates were transferred to a 37°C incubator for 15-20 min until hybrid-hydrogels polymerized. Constructs were allowed to rest overnight in 10% FBS in DMEM:F12 before being transferred to fresh wells. Media was replaced every 2-3 days for one week before dynamic stiffening.

3D images of model structure were generated by incubating epithelial cells and microspheres in CellTracker Green CMFDA (Fisher) during the 3-day aggregation procedure. After encapsulation with fibroblasts in hybrid-hydrogels, constructs were fixed in 4% PFA for 1 h, permeabilized with 0.2% Triton X-100, and blocked with 5% bovine serum albumin (BSA) in PBS. Constructs were incubated overnight at 4°C with a 1:100 dilution of anti-collagen antibody in 5% BSA/PBS (clone E8F4L, Cell Signaling Technology), washed three times for 15 min in PBS with 0.1% v/v Tween 20, and then incubated with ActinRed 555 ReadyProbes (2 drops/mL, Fisher) and AlexaFluor 647 anti-rabbit secondary (1:400, Fisher) for 1 h at room temperature before a final set of three washes. Imaging was performed on an inverted spinning disc confocal microscope (Leica 3I MARIANAS) using the 40x water objective. 3D projections were generated using the orthogonal views function in Fiji (ImageJ).

### Dynamic stiffening

A solution of ruthenium (0.47 mM, Sigma Aldrich) and sodium persulfate (SPS; 5 mM, Sigma Aldrich) was prepared in 1% FBS in DMEM:F12 and allowed to diffuse into 3D hydrogels for 2 h at 37°C until the solution had visibly permeated the full construct. 3D lung models were then transferred to PBS, exposed to 440 nm light for 5 min, placed back into 1% FBS in DMEM:F12, and incubated at 37°C.

### Reporter and viability imaging

3D lung models containing dual reporter Col1a1-GFP and αSMA-RFP fibroblasts were incubated with 10 µM Hoechst and 1 µM thiazole red in 1% FBS DMEM:F12 for 1 h before being transferred to fresh media in a glass-bottom plate (MatTek). Samples (N=4) were imaged on a Keyence BZ-X800 microscope using the 10x objective. Three fields of view with a 100-µm z-stack were acquired per sample. All image analysis was performed in Fiji (ImageJ). Viability was quantified using the Analyze Particles function to count the number of total (Hoechst^+^) and dead (thiazole red^+^) nuclei. Marker expression was quantified on merged red/green images to count the number of red (αSMA^+^), green (Col1a1^+^), and yellow (dual positive) cells.

### RNA extraction and sequencing

Two weeks post-stiffening, 3D lung models were removed from 1% FBS in DMEM:F12 and a 1.5-mm biopsy punch separated the visible epithelial cell aggregate from the fibroblast-laden encapsulating hybrid-hydrogel. The separated fractions of 4-6 3D lung models were pooled into 300 ul Trizol in a 1.5 ml tube, flash frozen in liquid nitrogen, and then ground using an RNase-free pestle. Samples were allowed to thaw and rest at room temperature for 5 min before addition of 100 µl BCP. Samples were mixed, rested for an additional 5 min, and then centrifuged at 12,000 xg for 15 min at 4°C to achieve phase separation. The aqueous layer was transferred to an equal volume of 100% ethanol and mixed before being transferred to an RNeasy Mini spin column (Qiagen). Columns were centrifuged for 1 min at 8,000 xg and sequentially washed with buffer RW1 (Qiagen) and two changes of buffer RPE (Qiagen), spinning at the same settings for each wash. Columns were then transferred to clean collection tubes and spun at 17,000 xg for 2 min to dry the membrane. RNA was eluted into clean tubes by adding 24 µl nuclease-free water to the column and spinning at 17,000 xg for 1 min. Library preparation was performed using the SMARTer® Stranded Total RNA-Seq Kit (Takara Bio) without fragmentation for first-strand cDNA synthesis. Sequencing was performed using a low-input ribo-depleted total RNA-seq protocol with 80M paired-end reads per sample.

### RNAseq data processing

Raw RNA sequencing data were assessed using FastQC. We employed Cutadapt^28^ for adapter trimming. Reads were aligned to the mouse reference genome (GRCm38) using STAR.^29^ Gene-level read counts were quantified from the aligned BAM files using featureCounts. A gene-level count matrix for all samples was used for downstream statistical analyses in R version 4.5.1. We removed lowly expressed genes, which had fewer than 10 counts in more than 50% of the samples, resulting in 10022 genes across 61 samples. Differential expression analysis was performed using DESeq2 package,^30^ which models gene counts with a negative binomial distribution. For each cell type, multiple pairwise contrasts between conditions were specified. Within each contrast, p-values were adjusted for multiple testing across genes using the Benjamini-Hochberg procedure.^31^ No additional correction was applied across contrasts. Differentially expressed genes were ranked by the adjusted p-values and absolute log2 fold change. The top ranked genes were used to generate heatmaps, bar plots, and volcano plot annotations. Differentially expressed genes were used for functional enrichment analysis to identify overrepresented biological processes and pathway using Gene Ontology database.

### Statistical methods

RNA extraction was performed on n=4-6 pooled constructs, and then sequencing run on N=5-7 separate pools. This sample number was calculated to achieve >80% power for detecting an effect size of 1.6 with a Gaussian distribution (1-sided t-test, p<0.05). For viability and reporter expression imaging experiments, a three-way ANOVA with Tukey’s multiple comparisons test was used to assess differences due to cell source, dECM source, and stiffness.

## Results and Discussion

### Generation of 3D lung models

3D distal lung models were generated by first mixing primary mouse lung epithelial cells with hydrogel microspheres. Both cells and spheres were magnetically labeled and then rotated together under a magnetic field for three days to allow the spheres to template the growth of epithelial cells into acinar-like structures. As previously published, this isolation technique results in an epithelial population that is approximately 85% surfactant protein C expressing AT2 cells, with an additional 13% expressing keratin 8 and 2% expressing podoplanin.^5^ On the third day, primary murine lung fibroblasts were isolated from either healthy or fibrotic mice to assess what cell-intrinsic phenotypes persist during *ex vivo* culture. Fibroblasts were combined with hybrid-hydrogel solution containing dECM from either healthy or fibrotic mouse lungs to investigate how ECM composition affects fibrotic development. The hybrid-hydrogels were comprised of a 10 mg/mol-1, 8-arm PEG-alpha methacrylate (PEGaMA) crosslinked by a 2:1:1 mixture of thiolated murine dECM, low-affinity MMP2-degradable peptide, and 34 kg/mol-1 PEG-dithiol.

The fibroblast-studded hybrid hydrogels were used to encapsulate epithelial cell aggregates in a model that mimics the 3D geometry of distal lung (Figure 1A). This is an advance over more commonly used dECM-alone hydrogels, which present difficulties in decoupling ECM composition and stiffness.^32^ More recent approaches to chemical functionalization and dynamic stiffening of dECM hydrogels has lent more tunability to these culture systems, but has yet to be employed in 3D co-culture models as here.^33,34^ Incorporation of the synthetic PEG-dithiol crosslinker was required to promote multi-week stability of the 3D lung models, particularly in conditions where fibroblast-mediated degradation of the biological crosslinkers was enhanced (Figure S1)

**Figure 1:**
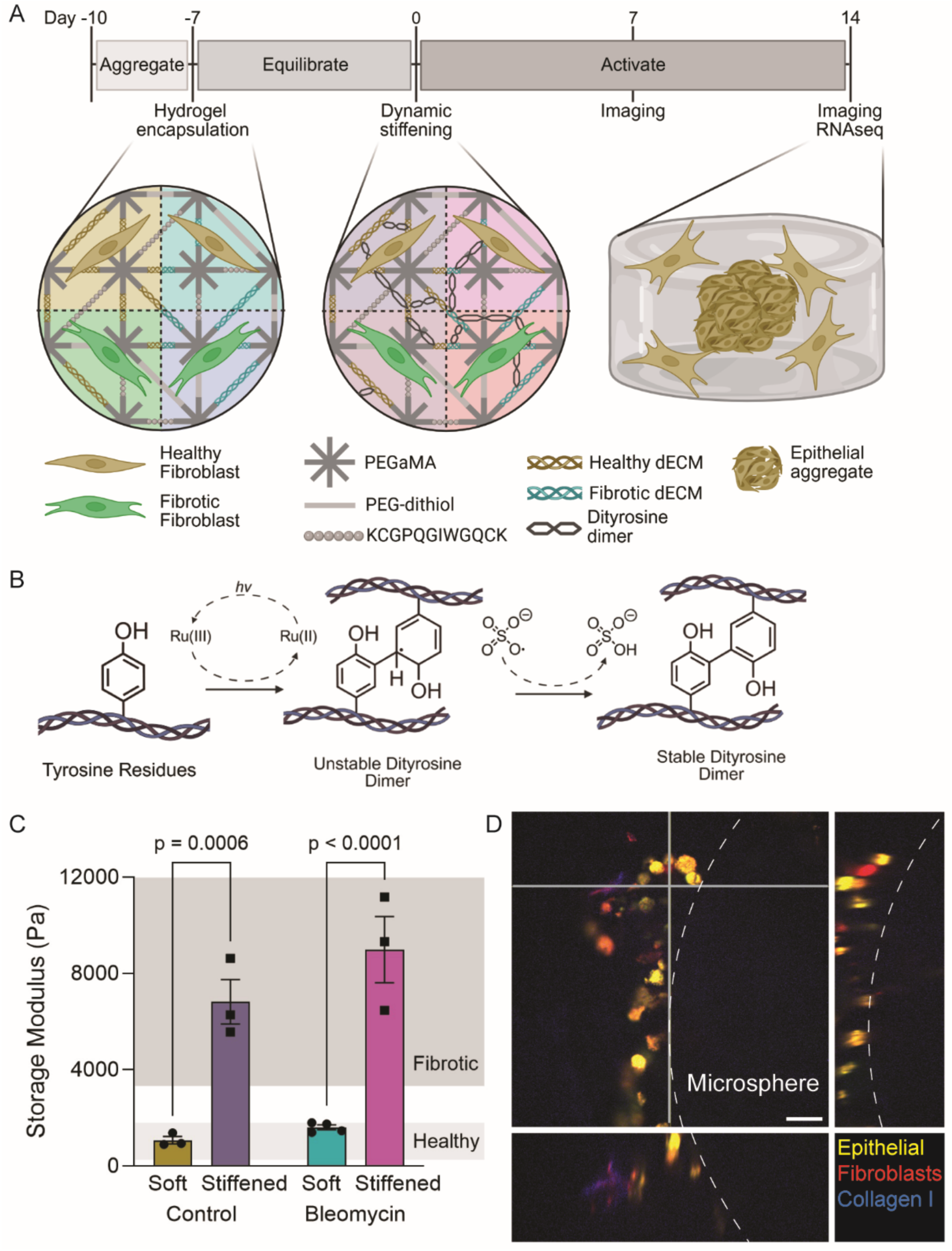
3D lung models mimic fibrotic inputs. A) Experimental timeline and overview of conditions. Epithelial cells were magnetically aggregated into acinar-like structures and then encapsulated with either healthy or fibrotic fibroblasts in hybrid-hydrogels containing either healthy or fibrotic dECM. After one week, hydrogels were either left soft or dynamically stiffened into the range of fibrotic tissue. B) Schematic of light (hv) initiated, ruthenium (Ru)/SPS mediated dityrosine dimerization stiffening reaction. C) Storage modulus of hybrid-hydrogels. N=3-4 batches D) 3D structure of lung models showing a region where dECM (collagen) and fibroblasts are near the epithelial cell layer. Scale = 20 µm

3D lung models were allowed to equilibrate for one week before dynamic stiffening to mimic the progressive tissue stiffening seen in lung fibrosis, with initially soft hydrogels (∼4 kPa) undergoing visible-light initiated dityrosine dimerization, resulting in 5-fold stiffening (Figure 1B, C). Published measurements of control vs IPF lung stiffness vary widely depending on species, measurement modality, and complicating factors including lung region, subject age, and subject sex.^35^ Studies have reported stiffness fold changes ranging from 2.5 to 18 as a result of fibrosis;^35^ our values were based on atomic force microscopy reading of human lung.^12^ Reactive oxygen species levels generated by the dynamic stiffening reaction remained below the *in vivo* threshold level across the course of the reaction (Figure S2), highlighting, as previously published, the cytocompatibility of this reaction.^36,37^ Viability and fibroblast phenotype were assessed at 1 and 2 weeks post-stiffening, with RNA isolation for genomics assessment at 2 weeks post-stiffening.

### Fibroblast phenotype imaging

Cellular viability and fibroblast marker expression were assessed across 8 conditions, comprising all the possible combinations of models containing fibroblasts from healthy or fibrotic lung, hybrid-hydrogels containing healthy or fibrotic dECM, and either remaining soft or undergoing dynamic stiffening (Table 1). This experimental design allowed us to decouple and investigate how cellular conditioning, matrix composition, and tissue stiffness contribute to fibrotic progression.

**Table 1:** Eight experimental conditions were assessed in imaging experiments. RNAseq was performed on conditions 1, 2, 3, 5, and 8.

| Condition | Fibroblast source | dECM source | Modulus |
| --- | --- | --- | --- |
| 1 | Healthy | Healthy | Soft |
| 2 | Fibrotic | Healthy | Soft |
| 3 | Healthy | Fibrotic | Soft |
| 4 | Fibrotic | Fibrotic | Soft |
| 5 | Healthy | Healthy | Stiffened |
| 6 | Fibrotic | Healthy | Stiffened |
| 7 | Healthy | Fibrotic | Stiffened |
| 8 | Fibrotic | Fibrotic | Stiffened |

Cell viability remained high across all conditions (Figure S3), confirming that hybrid-hydrogel polymerization and stiffening reactions were well-tolerated by cells. Fibroblast phenotype was assessed by expression of two endogenous markers: RFP under the control of the αSMA promoter and GFP under the control of the Col1a1 promoter (Figure 2A). Overall dynamics were quantified by the number of cells in the population expressing either marker, while fibroblast phenotype was represented by determining the percentage of cells that were either dual-positive or single positive for each marker. Interestingly, the strongest input affecting overall fibroblast activation marker expression was hydrogel stiffness. Stiffened samples all displayed around 30% marker expression one-week post-stiffening (Figure S4) and 60% by two weeks post-stiffening (Figure 2B), while soft samples showed approximately 20% marker expression after one week and ranged from 25-45% after two weeks. Both fibroblast and dECM source showed more modest impacts on marker expression, with inclusion of fibrotic dECM increasing activation marker expression around 1.6-fold inclusion of fibrotic fibroblasts around 1.8-fold, all after two weeks. The effects of stiffness on overall marker expression appeared to overcome differences induced by fibroblast and dECM source, as there were no notable differences between any of the stiffened conditions. Human IPF is characterized and driven by fibroblast heterogeneity; classic αSMA-expressing myofibroblasts appear to play strong roles on matrix remodeling via increased contractility, while ECM-synthesis is largely performed by differentially activated matrix fibroblasts, to name just two abundant populations.^38^ In our analysis of fibroblast phenotype, across all conditions the largest proportion of reporter-expressing fibroblasts were single-positive for Col1a1, followed by single-positive for αSMA, with the smallest proportion of fibroblasts expressing both markers (Figure 2C). Interestingly, in this analysis stiffness had a very limited effect, while inclusion of fibrotic dECM raised the proportion of dual-positive cells from 3% to 8%, and fibroblasts from fibrotic lung showed a high percentage of dual-positive cells in all conditions, around 25%. This is in line with data we have previously published using 2D culture on hybrid hydrogels, where stiffening caused around a 2-fold increase in activation, while fibrotic composition had an intermediate effect.^23^ Overall, these data suggest that environmental stiffness strongly impacts how many fibroblasts enter an activated state, while dECM composition or preconditioning in fibrotic lung still affect specific fibroblast phenotypes. To more closely examine cellular phenotypes in 3D models, we thus turned to transcriptomic analysis.

**Figure 2:**
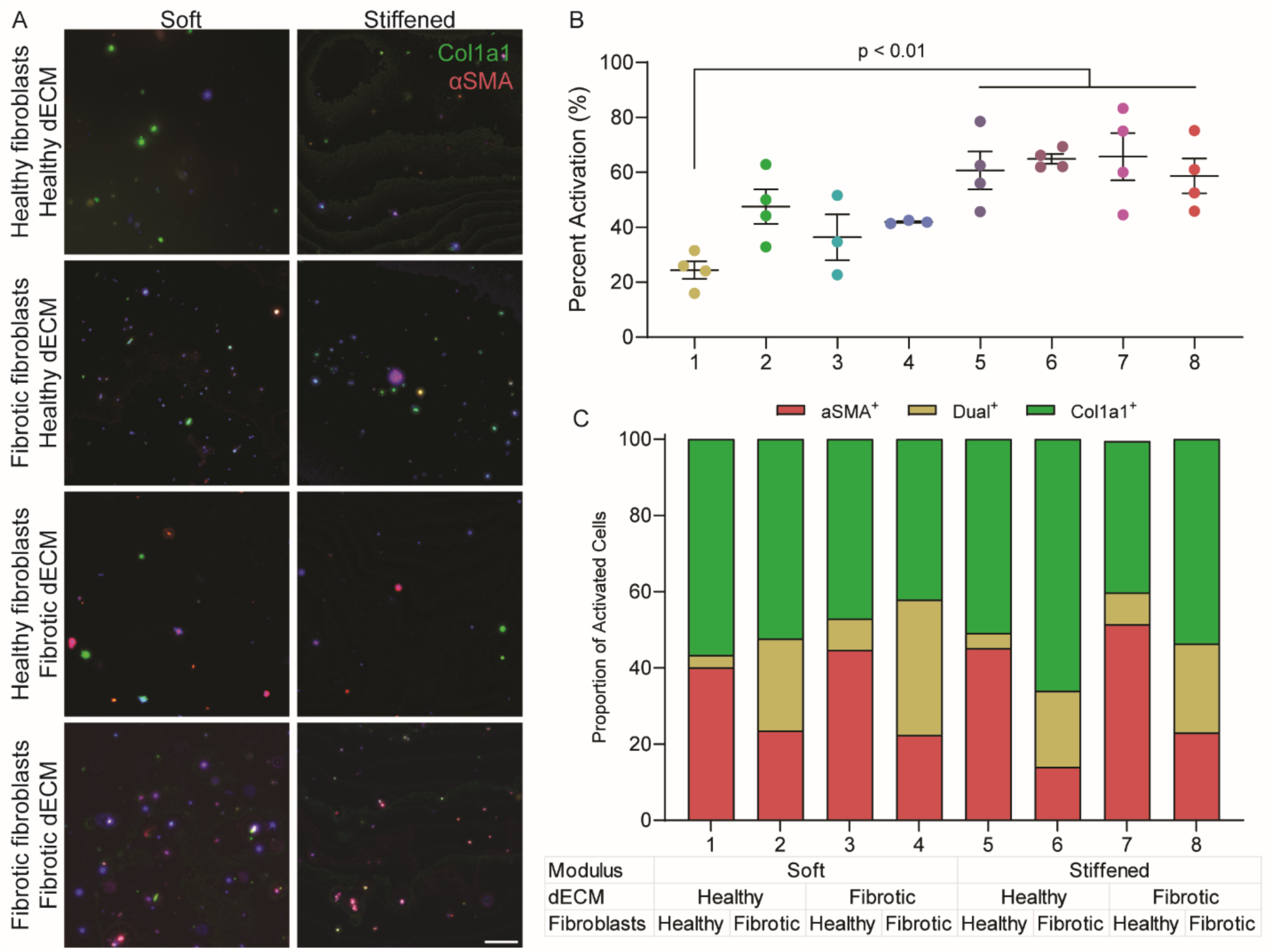
Fibrotic inputs uniquely contribute to overall fibroblast phenotype. A) Representative images of Col1a1-GFP/αSMA-RFP reporter fibroblast populations. Scale = 50 µm B) Quantification of fibroblast imaging showing the percentage of total cells that express either activation marker. N=3-4 constructs per condition, analysis by three-way ANOVA. C) Percentage of reporter-expressing cells that are single positive for either marker or dual positive.

### Transcriptomic analysis of 3D lung models

Five of the eight total conditions were selected by design-of-experiments statistical approach to assess in detail what genomic changes were induced by each experimental factor (fibroblast course, dECM source, and matrix stiffness) and interactions between these inputs. Condition 1 was designed to approximate healthy lung tissue and served as a reference condition, consisting of healthy murine fibroblasts encapsulated within a soft hybrid-hydrogel containing healthy lung dECM. Conditions 2, 3, and 5 were selected to isolate the effect of altering a single experimental factor while the remaining variables remained constant. In contrast, Condition 8 combined the fibrotic level of each input variable (fibrotic murine fibroblasts, fibrotic lung dECM, and increased matrix stiffness) and represented the hypothesized maximal fibrotic microenvironment. 3D lung models were microdissected generating central, epithelial cell-enriched and peripheral fibroblast-enriched regions for RNA extraction and genomic analysis.

Across both epithelial-enriched and fibroblast-enriched regions, stiffness exerted the strongest influence on cellular phenotype. Increasing matrix stiffness resulted in 2,407 differentially expressed genes (DEGs) in the epithelial-enriched regions and 3,051 in the fibroblast-enriched regions. In comparison, incorporation of fibrotic dECM into hybrid-hydrogels induced 343 DEGs in the epithelial-enriched regions and only 7 in the fibroblast-enriched regions. Likewise, inclusion of fibroblasts isolated from fibrotic lung showed 995 DEGs in the epithelial-enriched regions and 31 in the fibroblast-enriched regions (Figure 3A, B). Together these findings indicate that although fibrotic fibroblasts and fibrotic dECM each altered gene expression, matrix stiffening elicited a substantially broader transcriptional response in both cell populations. These results are consistent with the fibroblast imaging data, further supporting matrix stiffness as the dominant driver of cellular phenotype changes within this model.

**Figure 3:**
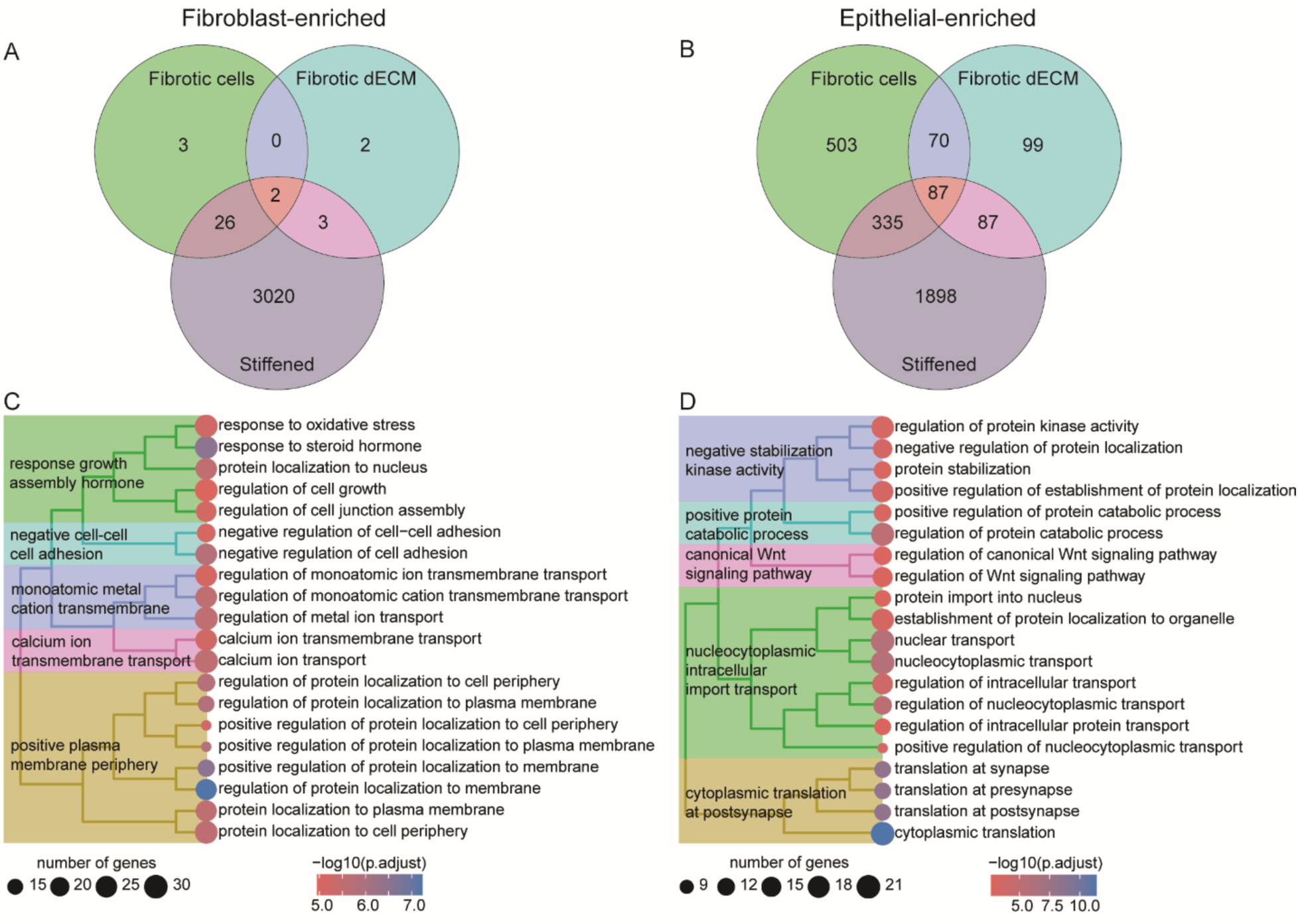
3D lung models recapitulate transcriptional programs associated with pulmonary fibrosis. Venn diagrams showing the number of differentially expressed genes (DEGs) in A) fibroblast-enriched regions and B) epithelial-enriched regions of 3D lung models attributable to incorporation of fibrotic fibroblasts, fibrotic dECM, and increased matrix stiffness. C-D) Gene ontology enrichment analysis of DEGs identified between the healthy reference condition (Condition 1) and the maximal fibrotic condition (Condition 8) in C) fibroblast-enriched regions and D) epithelial-enriched regions, highlighting biological processes altered in response to the combined fibrotic experimental factors. N=5-7 per condition.

Comparison of the healthy reference condition (Condition 1) with the maximal fibrotic condition (Condition 8) revealed transcriptional changes associated with pulmonary fibrosis in patient datasets.^39,40^ In the fibroblast-enriched regions, DEGs were enriched for pathways involved in oxidative stress,^9^ cellular growth,^10^ cell adhesion,^11^ protein localization to the plasma membrane,^11,41^ and transmembrane ion transport^41^ (Figure 3C). Collectively, these pathways are implicated in pathological fibroblast activation.^38^ Transmembrane transport is of particular interest, since these pathways link mechanosensing to the sustained metabolic and inflammatory gene expression pathways that characterize activated fibroblasts.^41^ A classic example of this link is the Piezo family of transmembrane ion channels, which open in response to mechanical stress, initiating calcium influx and leading to increased fibroblast activation and production of ECM mediators.^42,43^ These findings are consistent with our observation that matrix stiffness was the strongest driver of transcriptional changes across the engineered microenvironments. Although numerous 2D studies, including prior work from our group, have identified matrix stiffness as a key regulator of fibroblast activation,^14,16,23^ early studies in 3D hydrogels suggested that increasing matrix stiffness alone was insufficient to induce a profibrotic phenotype.^44,45^ These contrasting findings have largely been attributed to the inability of cells encapsulated within non-degradable or highly constrained hydrogels to spread and generate the traction forces required for mechanosensing. In contrast, our cell-degradable hybrid-hydrogels permitted fibroblast spreading and matrix remodeling before dynamic stiffening, enabling cells to establish mechanical interactions with the surrounding matrix.^5^ Under these conditions, increased matrix stiffness remained the dominant regulator of fibroblast transcriptional remodeling despite the simultaneous presence of fibrotic fibroblasts and diseased dECM, demonstrating that matrix mechanics continue to govern fibroblast activation in a physiologically relevant 3D microenvironment.

In the epithelial-enriched fraction, differentially expressed pathways included regulation of kinase activity, intracellular protein transport, cytoplasmic translation, protein localization, and Wnt signaling (Figure 3D). These pathways suggest broad remodeling of epithelial signaling and protein homeostasis in response to the fibrotic microenvironment. Aberrant Wnt signaling has been widely implicated in pulmonary fibrosis where it contributes to both epithelial dysfunction and fibroblast activation.^46–48^ Recent human single-cell and spatial transcriptomic studies similarly identified activation of Wnt, kinase, and injury-response pathways within epithelial populations localized to fibrotic niches, highlighting the importance of the surrounding microenvironment in directing epithelial cell state.^39,40^ The observation that these pathways emerge in our engineered 3D lung models suggests that these reductionist systems are sufficient to recapitulate key components of the transcriptional programs observed in human pulmonary fibrosis.

### Fibrotic fibroblasts

When 3D lung models containing fibroblasts isolated from fibrotic lungs (Condition 2) were compared to the healthy reference condition (Condition 1), the fibroblast-enriched regions showed a transcriptional program characteristic of fibroblast activation. Fibroblasts from healthy lungs overexpressed the uncharacterized developmental gene 4833420G17Rik^49^ as well as Sra1, which has been observed as an activated fibroblast marker in cardiac fibrosis.^50^ In contrast, fibroblasts from fibrotic lung strongly overexpressed Spp1 (osteopontin) (Figure 4A), the most highly upregulated gene identified in this comparison. Spp1 is one of the most consistently upregulated genes in human IPF, and is highly expressed within fibrotic foci particularly by pro-fibrotic macrophages but also by activated fibroblasts.^39,51^ In fibroblasts, Spp1 promotes integrin-mediated signaling through pathways including PI3K/AKT/mTOR, regulating proliferation, adhesion, and ECM remodeling,^52–54^ target pathways of the anti-fibrotic drugs pirfenidone and nintedanib.^55^ The persistence of Spp1 expression following encapsulation suggests that fibroblasts isolated from fibrotic lungs retain key features of the activated *in vivo* phenotype even within the engineered 3D lung model microenvironment. These opposing activation markers could suggest that even where healthy fibroblasts undergo activation *in vitro*, the cells may acquire different fibrotic phenotypes than those that are activated *in vivo*.

**Figure 4:**
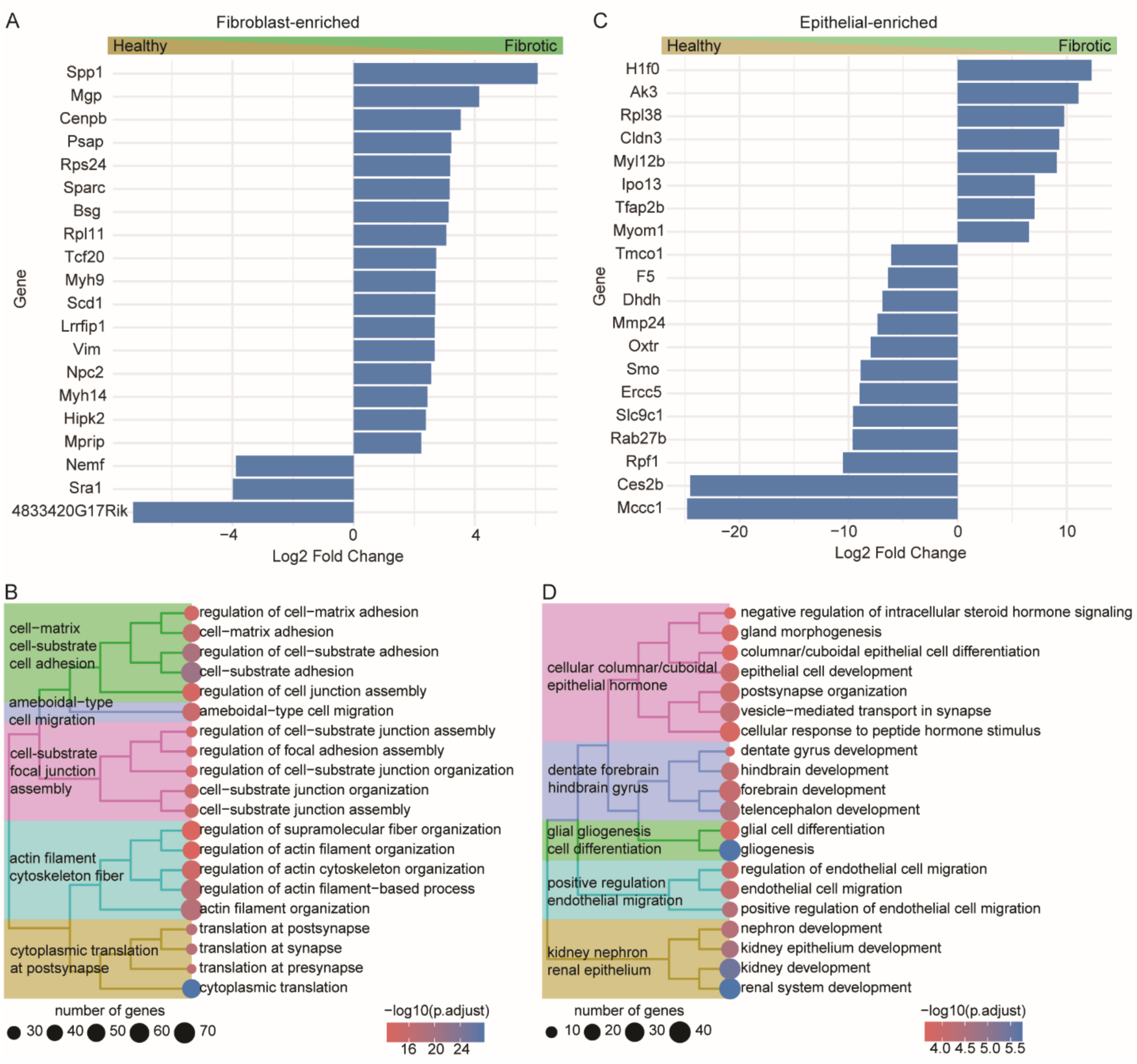
Fibroblast origin influences transcriptional programs in both fibroblast- and epithelial-enriched regions of 3D lung models. A) Top differentially expressed genes and B) gene network enrichment analysis in the fibroblast-enriched regions following incorporation of fibroblasts isolated from fibrotic (Condition 2) versus healthy (Condition 1) lungs. C) Top differentially expressed genes and D) gene networks in the epithelial-enriched regions, demonstrating that fibrotic fibroblasts induce transcriptional changes in epithelial developmental and differentiation pathways. N=5-7 per condition.

Consistent with these gene-level differences, pathway analysis of fibroblast-enriched regions (Figure 4B) showed enrichment of biological processes related to cellular adhesion, as represented by genes like Kdr and Plec, migration (Kdr, Myh9), cytoskeletal dynamics (Cnn2, Vim), and overall translation (Rps and Rpl family members). These pathways are characteristic of activated fibroblasts and closely parallel transcriptional programs identified in human IPF, where changes in cell-matrix adhesion and cytoskeletal remodeling support persistent fibroblast activation.^56–58^ Together, these findings suggest that fibroblasts isolated from fibrotic lungs retain a stable profibrotic transcriptional state that persists *in vitro*.

Inclusion of fibroblasts from fibrotic lungs also impacted gene expression within the epithelial-enriched regions. Two genes were notably upregulated in models containing healthy fibroblasts: Ces2d and Mccc1 (Figure 4C), both of which may have anti-fibrotic activity. Ces2 has predominantly been studied in the liver, where it protects against obesity-related inflammation and fibrosis.^59,60^ Mccc1 has not been widely studied in the context of fibrosis; however, genomic analysis has demonstrated reduced expression in IPF and a negative association with TGFβ signaling, suggesting a potential anti-fibrotic role.^61^ Pathway analysis showed enrichment of biological processes related to epithelial cell development, epithelial cell differentiation, and epithelial cell migration (Figure 4B). Although several enriched terms reference developmental processes in other organs, it is well known that there are overlaps between lung developmental pathways and the dysregulated epithelial repair programs that are activated during IPF.^62^ Many of the genes involved in these developmental pathways also participate in canonical fibrotic signaling cascades, including TGFβ and Sonic Hedgehog (Shh) signaling.^63^ Human single-cell and spatial transcriptomic studies similarly reveal that epithelial populations adjacent to fibrotic fibroblasts exhibit dysregulated developmental and differentiation programs, supporting the concept that crosstalk with fibrotic fibroblasts may impact epithelial cell differentiation and acquisition of pathological phenotypes.^3,39,64^

### Fibrotic dECM

Hybrid-hydrogels containing dECM from fibrotic lungs (Condition 3) induced relatively modest but biologically relevant differences (relative to Condition 1) in the cellular phenotype of both fibroblasts (Figure 5A) and epithelial cells (Figure 5B). In the fibroblast-enriched regions, the small number of upregulated genes converged on pathways involved in development, tRNA methylation, and the Jnk signaling cascade (Figure 5C). RNA methylation as a driver of fibrosis is still understudied, but has been implicated in the pathogenesis of pulmonary, cardiac, and hepatic fibrosis.^65–67^ In fibroblasts specifically, tRNA methytrasferase activity has been observed to promote proliferation, activation, and migration.^68,69^ Among the enriched pathways, JNK signaling, meanwhile, is well-known to promote fibroblast activation downstream of ECM remodeling and integrin signaling while intersecting with multiple canonical profibrotic pathways, including TGFβ, JAK/STAT, Wnt, and PI3-kinase pathways.^70–73^ These results suggest that while fibrotic dECM alone induces fewer transcriptional changes, it selectively activates signaling pathways known to regulate fibroblast responses to ECM remodeling and shifts in these pathways, therefore, show that exposure to fibrotic dECM in hybrid-hydrogel environments can impact fibroblast activation phenotype.

**Figure 5:**
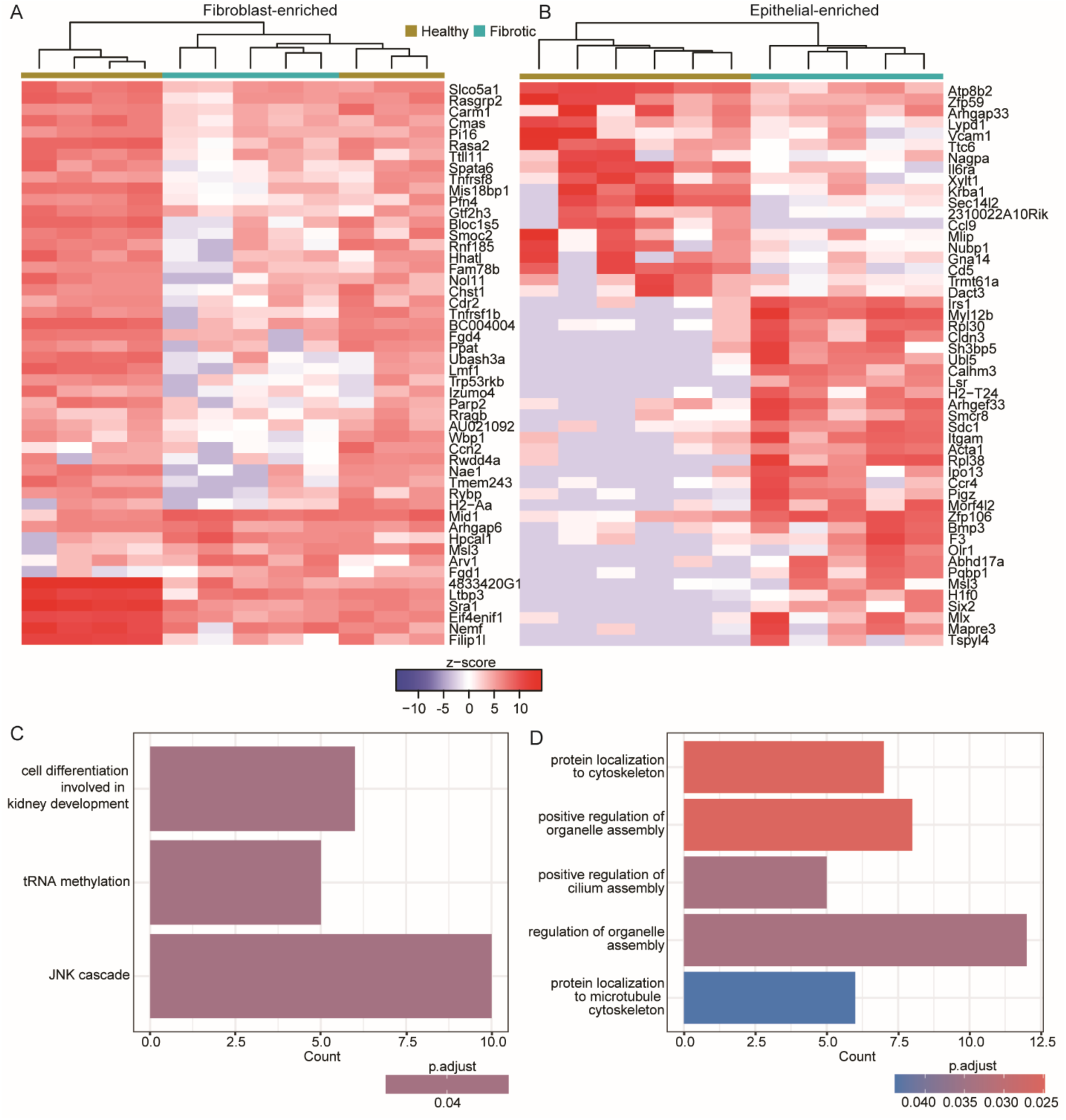
Fibrotic dECM induces selective transcriptional changes in 3D lung models. Heatmaps of high-fold change genes in A) the fibroblast-enriched and B) the epithelial-enriched regions comparing incorporation of fibrotic dECM (Condition 3) to the healthy reference condition (Condition 1). Differentially regulated gene networks in C) the fibroblast-enriched and D) the epithelial-enriched regions, showing enrichment of stress-response, developmental, and intracellular organization pathways following exposure to fibrotic dECM. N=5-7 per condition.

In the epithelial-enriched regions, fibrotic dECM altered pathways involved in intracellular organization, including protein localization, cytoskeletal dynamics, and organelle and cilium assembly (Figure 5D). Changes in cytoskeletal organization are a hallmark of epithelial-to-mesenchymal transition, a program that has been associated with response to pathological ECM remodeling and acquisition of pro-fibrotic phenotypes.^74,75^ Relatedly, cilium assembly relies on cytoskeletal involvement, and suggest the development of more motile cellular phenotypes, which has been studied in the context of fibroblast activation,^76,77^ but may also be involved in bronchiolization during fibrosis, as epithelial cells from airways travel into distal lung.^78,79^ Primary cilia also function as important signaling organelles that coordinate Shh, Wnt, and other developmental pathways in pulmonary fibrosis.^80,81^ Altered cilium assembly may reflect changes in epithelial signaling and differentiation associated with dysregulated repair. Together, these results indicate that while the number of genes altered by ECM composition alone is smaller than transcriptional changes in response to fibroblast origin or matrix stiffening, the affected pathways induced by hybrid-hydrogels are deeply relevant to epithelial and fibroblast behaviors that contribute to fibrotic disease progression.

### Stiffness

Dynamic stiffening (Condition 5) induced the most profound transcriptional changes of the three variables investigated (relative to Condition 1). In the fibroblast-enriched regions, glutathione s-transferase (GST) family members including Gstp1 and Gsta1 were overexpressed in soft samples (Figure 6A). These genes contribute to cellular antioxidant defenses and reduction of GST activity, including by genetic polymorphisms, can increase risk of fibrotic responses.^82^ In contrast, stiffened hybrid-hydrogels promoted overexpression of profibrotic genes including Tns1, a regulator of focal adhesions, which is implicated in fibroblast activation and ECM production,^83^ and Card10, which promotes collagen deposition and inflammation in mouse models of fibrosis.^84^ Beyond individual genes, dynamic stiffening induced coordinated enrichment of pathways governing cell-matrix interactions, cytoskeletal dynamics, GTPase signal transduction, response to mechanical stimulus, and TGFβ signaling (Figure 6B). Together, these pathways comprise a canonical mechanotransduction program through which matrix stiffness is translated into intracellular signaling via the action of GTPases like Rho and changes to the actin cytoskeleton, ultimately driving alterations in cellular proliferation, migration, contractility, and ECM remodeling.^16,85,86^ One important relationship between TGFβ signaling and environmental stiffness is in the transformation from latent to active protein, which can be mediated by conformational changes induced by cellular contraction of the ECM.^87^ Because our hybrid-hydrogels incorporate native lung dECM, which we have previously shown retains a variety of extracellular growth factors,^24^ dynamic stiffening of our hybrid-hydrogels may enhance activation of matrix-bound TGFβ while simultaneously priming fibroblasts to respond through mechanosensitive pathways that converge on TGFβ/SMAD signaling, an interaction well known to be at the heart of fibrotic progression.^57^ Collectively, these results demonstrate that dynamic stiffening recapitulates the coordinated mechanobiological pathways that regulate fibroblast activation during pulmonary fibrosis.^58^

**Figure 6:**
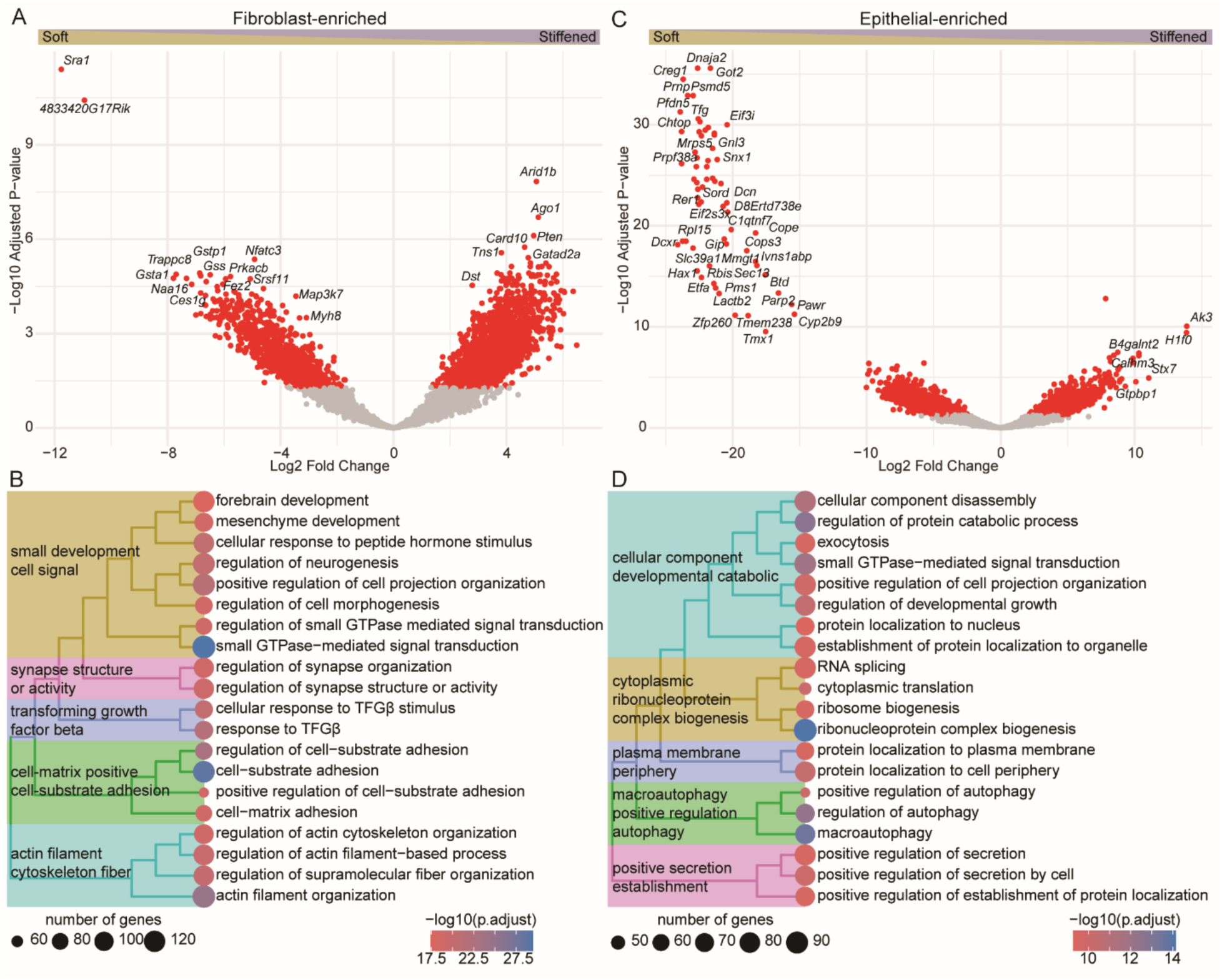
Dynamic stiffening drives widespread transcriptional changes in 3D lung models. A) Volcano plot of differentially expressed genes and B) top differentially regulated gene networks in fibroblast-enriched regions following dynamic stiffening (Condition 5 relative to Condition 1), showing enrichment of pathways associated with mechanotransduction, cell-matrix interactions, cytoskeletal organization, and TGFβ signaling. C) Volcano plot of differentially expressed genes and D) top differentially regulated gene networks in the epithelial-enriched regions, highlighting extensive changes in pathways related to protein homeostasis and autophagy in response to increased matrix stiffness. N=5-7 per condition.

Dynamic stiffening also notably produced extensive transcriptional changes in the epithelial-enriched regions (Figure 6C). A large cluster of genes, including Creg1, which has been studied for its protective role against cardiac fibrosis via activation of autophagy,^88^ was more highly expressed in soft hydrogels. In stiffened hydrogels, meanwhile, overexpressed genes included pro-fibrotic factors like H1f0, a histone associated with cellular stress responses, fibroblast activation, and fibrotic progression.^89^ Analysis demonstrated enrichment of pathways involved in protein localization, translation, secretion, autophagy, and intracellular organelle organization (Figure 6D), indicating widespread remodeling of epithelial protein homeostasis and secretory function in response to increased matrix stiffness. Autophagy has been reported to be overall downregulated during the progression of IPF, in both fibroblasts and epithelial cells.^90–92^ Treatments that induce autophagy also reduce fibrotic responses in experimental models, suggesting an overall protective role for autophagy in resolving cellular dysfunction associated with IPF.^92,93^ Although the relationship between environmental stiffness and autophagy remains context dependent, most previous studies have reported that stiffness can increase autophagy in smooth muscle cells,^94^ pancreatic cancer cells,^95^ and fibroblasts^96^ while decreasing autophagy in endothelial cells.^94,97^ Critically, the majority of these studies were performed in 2D culture.^17^ Our findings show that dynamic stiffening alone is sufficient to alter autophagy-associated transcriptional programs within a cell-degradable 3D lung model, highlighting the importance of matrix mechanics in regulating epithelial cell homeostasis and the importance of further study of these interactions in physiologically relevant biomaterial systems. These results together with the factorial transcriptomic analysis findings, reveal that matrix stiffness is the dominant regulator of both fibroblast and epithelial transcriptional programs, exceeding the effects of fibrotic fibroblast origin or ECM composition alone.

### Epithelial signatures in murine and human data

Since we observed the greatest changes in transcriptomic profiling within the epithelial-enriched fraction, we queried some of our top hit genes against the IPF cell atlas, a multi-study database of human single-cell sequencing data.^98–100^ Our data revealed 23 genes that were conserved across all comparisons, many of which were upregulated in human IPF-associated epithelial cells, particularly transitional and aberrant basaloid cells (ABECs). Of particular interest was LBH, a TGF-β induced transcriptional co-factor that has been implicated in cardiac,^101^ liver,^102^ and renal fibrosis,^103^ and is expressed in human ABECs 10-12 fold higher than in AT2 cells. Our models also identified IRS1, which is expressed in ABECs at levels 3-5 fold higher than in AT2 cells and links insulin signaling and the PI3 kinase cascade in liver fibrosis.^104^ These results suggest, in line with our previously published results,^5^ that even epithelial cells from healthy mice can acquire pathogenic phenotypes in biomaterial-based 3D lung models, creating a valuable platform for the study of how epithelial cell biology contributes to both the initiation and progression of disease.

## Conclusions

Fibrotic lungs are characterized by changes in dECM composition, lung mechanics, and cellular activation, which are tightly interconnected, making it difficult to determine the individual contributions of each factor to disease progression. The present study experimentally decoupled these hallmark features of the fibrotic niche by combining dynamically stiffening hybrid-hydrogels containing disease-specific dECM with primary fibroblasts from healthy or fibrotic lungs within 3D lung models to investigate the relative contribution of these factors to fibrotic cellular phenotypes. Although all three experimental factors influenced cellular phenotype, matrix stiffening emerged as the strongest regulator of transcriptional remodeling, producing the largest number of DEGs and activating canonical mechanobiology pathways regulating cell-matrix interactions, cytoskeletal remodeling, TGFβ signaling, and epithelial protein homeostasis. In contrast, fibrotic dECM composition elicited a less pronounced transcriptional response centered on stress-responsive pathways, while fibroblasts isolated from fibrotic lungs retained a profibrotic transcriptional program that altered epithelial cell developmental and differentiation pathways. Collectively, these results demonstrate that biomechanical, biochemical, and cell-intrinsic cues each contribute to distinct aspects of fibrotic biology, while tissue mechanics remain the primary driver of transcriptional responses within engineered 3D lung microenvironments.

Across multiple conditions, hybrid-hydrogel based 3D lung models recapitulated key transcriptional programs associated with human pulmonary fibrosis. Consistently differentially regulated pathways included those involved in plasma membrane dynamics and secretion, development, and cytoskeleton and adhesion. In line with these results, certain genes emerged as differentially regulated across all conditions, including Sra1 and 4833420G17Rik in the fibroblast-enriched regions, and Dpp4, Usp5, and Itgam in epithelial-enriched regions. Among these, Dpp4 is particularly notable because the DPP4 serine peptidase has been observed on both alveolar epithelial type II cells and fibroblasts, where it appears to promote fibrotic progression via TGFβ signaling.^105,106^ Usp5 encodes for a deubiquitinase that has been implicated in the progression of lung cancer,^107^ and more recently in pathological epithelial differentiation in the context of fibrosis through regulation of NF-κB and hedgehog signaling.^108^ Itgam encodes for an integrin alpha subunit most commonly observed on macrophages/monocytes and involved in cellular adhesion, migration, and polarization.^109–111^ The repeated identification of these pathways and genes across various independent conditions further illustrates that engineered 3D lung models recapitulate molecular features of human pulmonary fibrosis.

One limitation of the current study is the microdissection method used to isolate fibroblast- and epithelial-enriched regions. The epithelial-enriched regions likely contained a subset of fibroblasts that were adjacent to the epithelial cell aggregates, potentially representing the most highly activated fibroblast population due to exposure to epithelial-cell derived activation factors. This cellular overlap may partially explain the overall larger number of DEGs within the epithelial-enriched regions across comparisons. Consistent with this interpretation, several notable genes identified within the epithelial-enriched regions, including Mccc1,^112^ H1f0,^89^ Creg1,^113^ and Dpp4,^106^ have previously been reported to be expressed by fibroblasts. Nevertheless, these data still support the relevance of 3D lung models as a platform for studying cell-level fibrotic phenotypes, and the power of hybrid-hydrogels as a unique platform to decouple tissue stiffness and ECM composition. Regional profiling distinguished biologically meaningful transcriptional responses within fibroblast-enriched and epithelial-enriched regions and revealed extensive epithelial-fibroblast crosstalk that would not have been captured using bulk transcriptomic approaches.

More broadly, this work establishes dynamically tunable hybrid-hydrogels as a versatile biomaterial platform for decoupling and investigating the independent and combined contributions of matrix mechanics, ECM composition, and cellular conditioning to pulmonary fibrosis. Future studies incorporating human cells and dECM will enable investigation of clinically relevant questions in IPF, including therapeutic responses, patient-specific disease heterogeneity and sex-dependent differences in fibrosis. Biomaterials capable of both dynamic stiffening and softening could facilitate studies of fibrosis resolution, while 3D models containing immune and/or vascular cells could be employed to interrogate increasingly complex multicellular fibrotic niches. Together, these findings establish hybrid-hydrogel-based 3D lung models as a powerful platform for mechanistic studies of pulmonary fibrosis and for the development and pre-clinical evaluation of next-generation anti-fibrotic therapies.

## Acknowledgements and Funding

RNAseq troubleshooting and performance was guided by Dr Schuyler Lee, associate director of the CU Anschutz Genomics Shared Resource. 3D imaging was performed in the Advanced Light Microscopy Core facility of the NeuroTechnology Center at the University of Colorado Anschutz, which is supported in part by the National Institute of Diabetes and Digestive and Kidney Diseases (P30 DK116073). Additional funding was provided by the National Heart, Lung, and Blood Institute HL153096 (RB, MCM, DE, CMM), National Science Foundation 1941401 (MCM, CMM), National Center for Advancing Translational Sciences UM1 TR004399 (TV), Public Health Service Grants HL140595 and HL162607 (DWHR), and U.S. Department of Veteran’s Affairs (VA) Merit Award BX003471 (DWHR). Contents are the authors’ sole responsibility and do not necessarily represent official NIH views.

## CRediT Authorship Statement

RB: Data curation, Formal analysis, Investigation, Methodology, Visualization, Writing - Original Draft, Writing - Review & Editing; MM: Investigation, Methodology; TV: Data curation, Formal analysis, Visualization, Writing - Review & Editing; DE: Investigation, Writing - Review & Editing; DR: Methodology, Writing - Review & Editing; CM: Conceptualization, Data curation, Funding acquisition, Project administration, Resources, Supervision, Visualization, Writing - Original Draft, Writing - Review & Editing

## Data Availability Statement

The data that support the findings of this study are openly available in Mendeley Data at doi: 10.17632/2hmgxs576g.2. Transcriptomic data can be found in the NCBI Gene Expression Omnibus (accession number to be assigned).

