## Supplemental Information for "Dynamic hybrid-hydrogel lung models decouple matrix composition, stiffness, and fibroblast conditioning to define distinct drivers of fibrotic progression"

**Supplemental Methods**

*Degradation analysis*

Hydrogels were fabricated by crosslinking 8-arm PEGαMA with crosslinkers mixtures containing 50% functionalized dECM and then various combinations of a high affinity MMP9-degradable peptide (Ac-GCRD-VPLSLYSG-DRCG-NH2), low affinity MMP2-degradable peptide (KCGPQGIWGQCK), and non-degradable PEG-dithiol. Fibroblasts were encapsulated in hydrogels at a concentration of 62,500 cells/15 µl and cultured for 28 days. In half the samples, dynamic stiffening was performed on day 7. The number of constructs was counted at each media change (every 48 hours) to track cell-mediated degradation of the hydrogel.

*Amplex red assay*

Amplex red detection of hydrogen peroxide was performed according to manufacturer direction (Fisher A22188). 1% serum media was supplemented with 0.47 mM ruthenium and 5 mM SPS and then rested for 4 hours at 37° to recapitulate the timeline of hydrogel swelling. Some media was taken for analysis, while the rest was exposed to 450 nm for 5 minutes as in dynamic stiffening, and then assessed immediately, at 30 minutes, 1 hour, and one day post-light to determine levels and persistence of ROS as a result of stiffening chemistry. Media samples and a hydrogen peroxide standard curve were reacted with 100 µM Amplex Red reagent and 0.2 U/mL horseradish peroxidase and read on a fluorescence plate reader at excitation 55 nm/emission 590 nm.

**Supplemental Results**

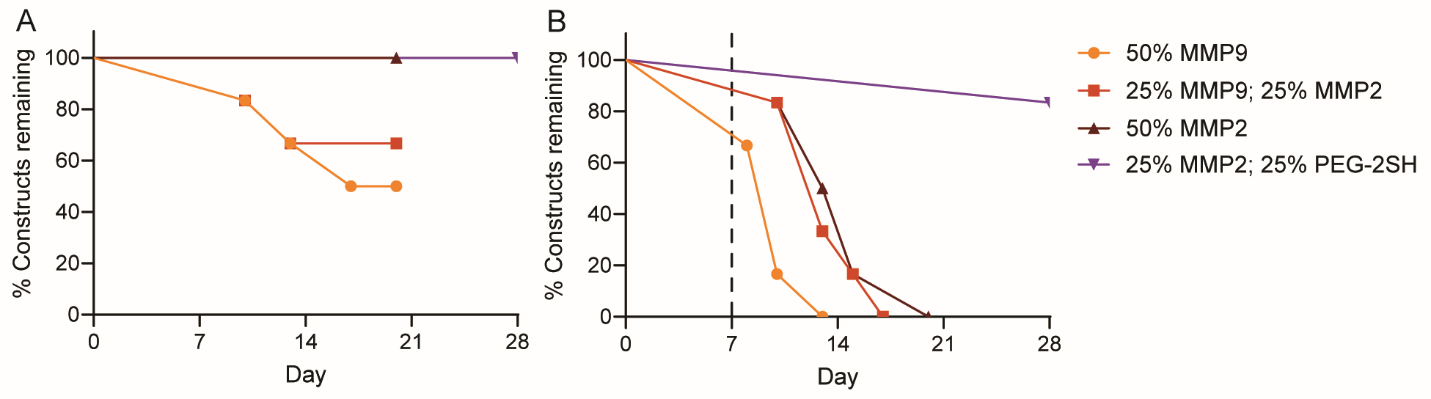

**Figure S1.** Degradation analysis of hydrogels with various crosslinker composition, either left A) soft or B) dynamically stiffened shows that 25% PEG-dithiol is required to prevent total enzymatic degradation of the constructs by three weeks.

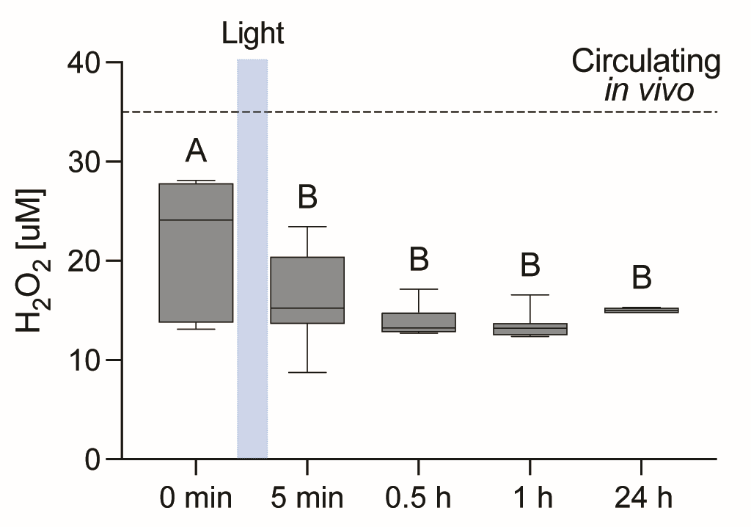

**Figure S2.** ROS levels in media containing SPS and Ruthenium for stiffening remain below the level detected *in vivo* across the time scale of cell exposure.

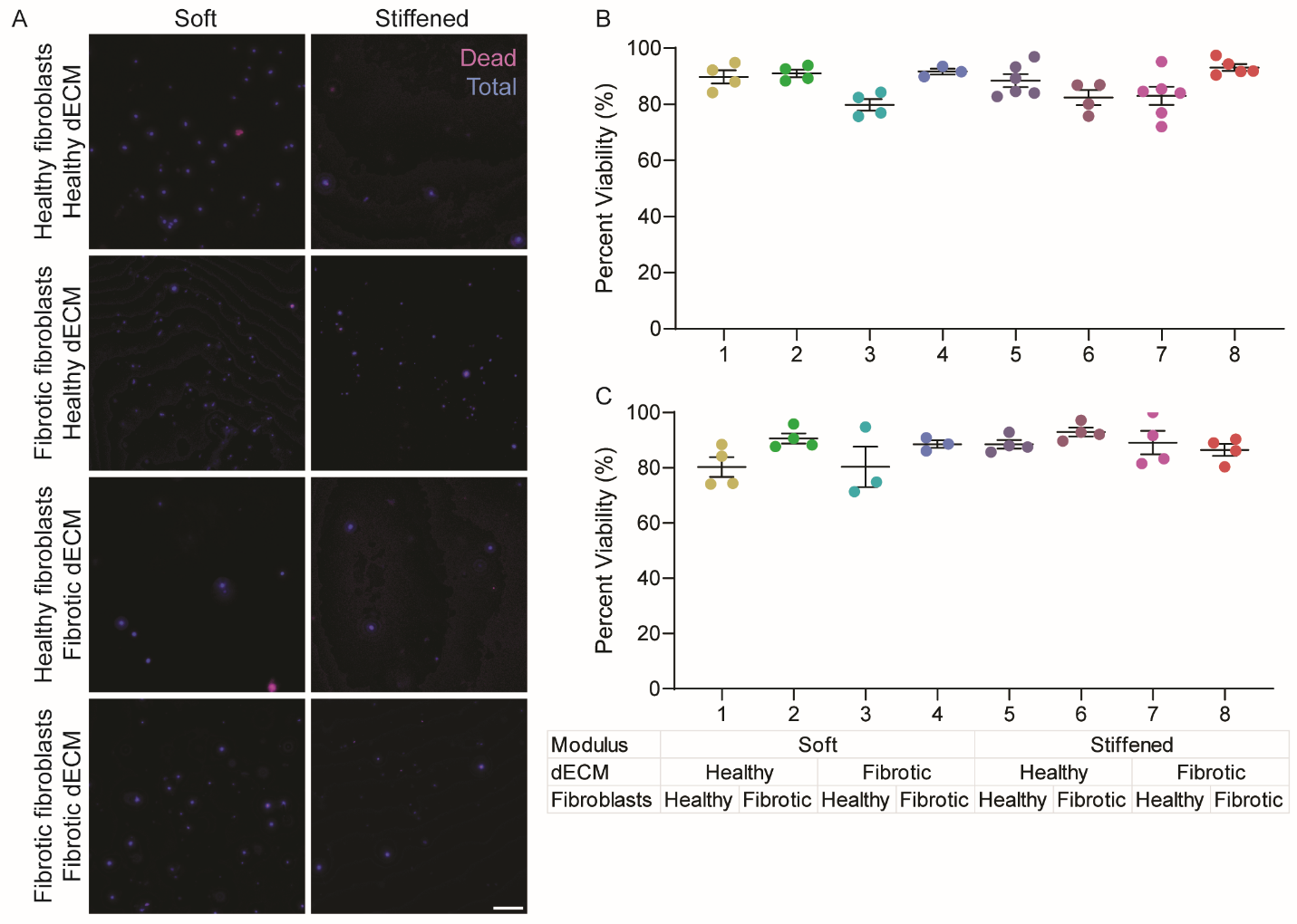
**Figure S2.** Cell viability in 3D lung models. A) Representative live-dead staining images from two weeks post-stiffening. Scale = 100 µm. Viability quantification at B) one week and C) two weeks post-stiffening.

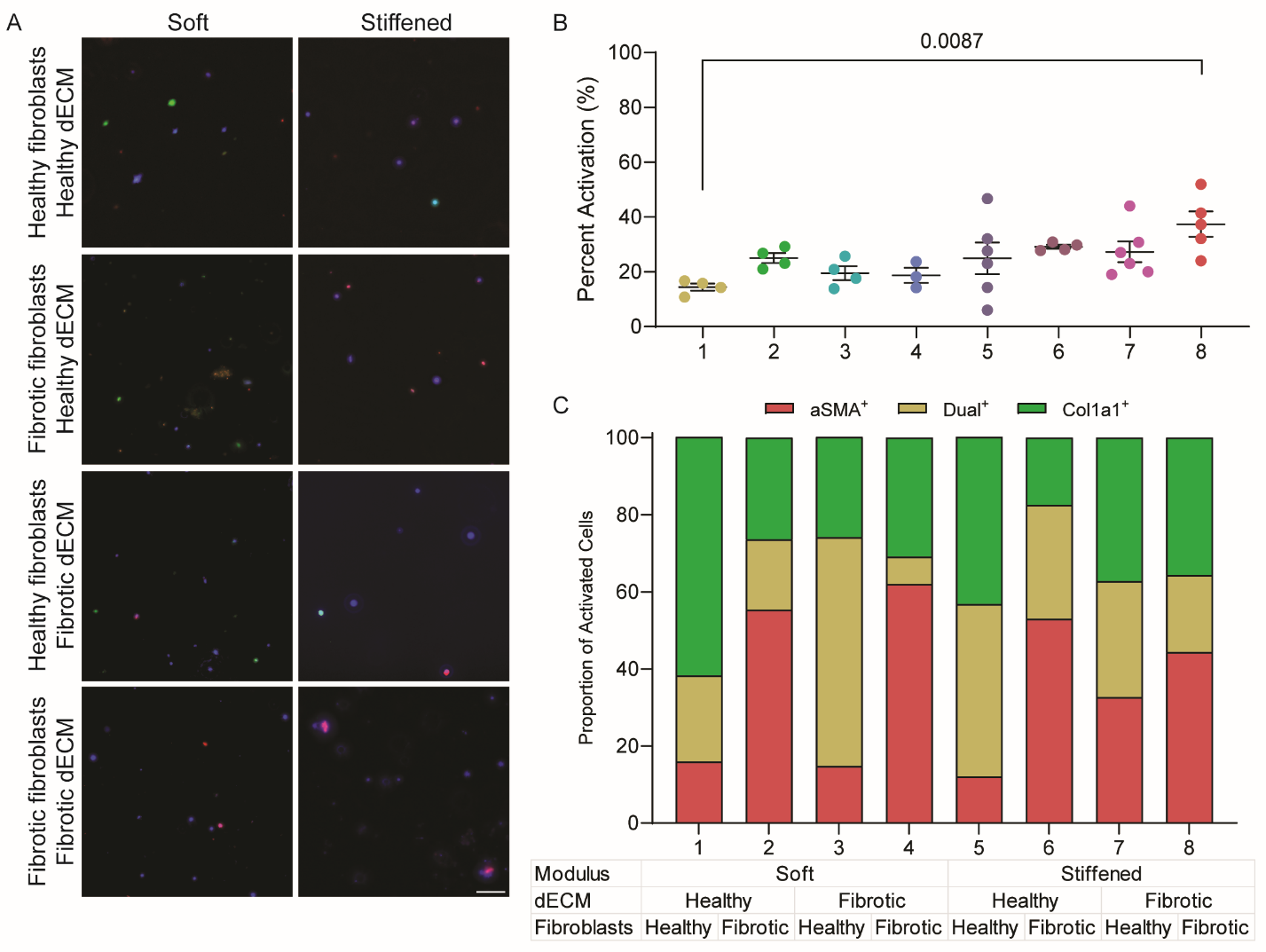

**Figure S3.** Fibroblast activation one-week post-stiffening. A) Representative images of dual-reporter fibroblast populations. Scale = 100 µm B) Quantification of overall phenotype by percentage of cells expressing either marker. C) Quantification of marker proportion across conditions.
